# From genome wide SNPs to genomic islands of differentiation: the quest for species diagnostic markers in two scleractinian corals, *Pocillopora* and *Porites*

**DOI:** 10.1101/2022.10.21.513203

**Authors:** Romane Deshuraud, Alexandre Ottaviani, Julie Poulain, Marine Leprêtre, Odette Beluche, Eric Mahieu, Sandrine Lebled, Caroline Belser, Alice Rouan, Clementine Moulin, Emilie Boissin, Guillaume Bourdin, Guillaume Iwankow, Sarah Romac, Sylvain Agostini, Bernard Banaigs, Emmanuel Boss, Chris Bowler, Colomban de Vargas, Eric Douville, Michel Flores, Paola Furla, Pierre Galand, Fabien Lombard, Stéphane Pesant, Stéphanie Reynaud, Matthew B Sullivan, Shinichi Sunagawa, Olivier Thomas, Romain Troublé, Rebecca Vega Thurber, Christian R. Voolstra, Patrick Wincker, Didier Zoccola, Serge Planes, Denis Allemand, Eric Gilson, Didier Forcioli

## Abstract

Coral reefs are of paramount importance in marine ecosystems, where they provide support for a large part of the biodiversity. Being quite sensitive to global changes, they are therefore the prime targets for biodiversity conservation policies. However, such conservation goals require accurate species identification, which are notoriously difficult to get in these highly morphologically variable organisms, rich in cryptic species. There is an acute need for easy-to-use and resolutive species diagnostic molecular markers. The present study builds on the huge sequencing effort developed during the TARA Pacific expedition to develop a genotyping strategy to assign coral samples to the correct species within two coral genera (*Porites* and *Pocillopora*). For this purpose, we developed a technique that we called “Divergent Fragment” based on the sequencing of a less than 2kb long diagnostic genomic fragment determined from the metagenomic data of a subset of the corals collected. This method has proven to be rapid, resolvable and cost-effective. Sequencing of PCR fragments nested along the species diagnostic fragment allowed us to assign 232 individuals of the genus *Pocillopora* and 247 individuals of the genus *Porites* to previously identified independent genetic lineages (*i*.*e*. species). This genotyping method will allow to fully analyze the coral samples collected across the Pacific during the Tara Pacific expedition and opens technological perspectives in the field of population genomics-guided conservation.

## Introduction

Coral reefs are natural structures which, although covering 0.2% of the ocean surface, host more than 25% of marine biodiversity (Chen et al. 2015). They therefore constitute essential marine ecosystems, providing ecological niches for many species. However, they are threatened by the effects of global changes, in particular the warming and acidification of the oceans (Masson-Delmotte et al. 2018).

In this context, the main objective of the Tara Pacific expedition is to analyze the complexity of coral reefs and their modifications in the face of climate change, as well as anthropic pressures (Planes et al. 2019). Thus, a multi-level multi-species dataset of the coral reef ecosystem across the Pacific Ocean is under construction thanks to the samples taken during the Tara schooner campaign which took place from 2016 to 2018 over more than 100,000 km and visited 32 islands over the entire Pacific transect. Three genera of corals were sampled, two scleractinian corals *Pocillopora* and *Porites*, as well as the hydrocoral *Millepora*. These coral samples are analyzed by multi-omics (metabarcoding, metagenomics, metatranscriptomics and metabolomics) and the quantification of molecular phenotypes (telomere DNA length, stress biomarkers). In addition, these samples are associated with a set of environmental physicochemical data recorded on site at the time of sampling and inferred from satellite and other data, including historical climatic parameters (Planes et al. 2019) (Lombard et al. 2022). This warehouse of data should provide a better understanding of the complexity of reef-building corals and their responses to climate change (Planes et al. 2019).

However, the phenotypic response to climate change can only be properly analyzed if the studied individuals are assigned to species, as the species, viewed as a shared gene pool, is what constitutes the adaptive toolbox (see for example Boulay *et al*., 2014). The initial set of analyses on *Pocillopora* and *Porites* from the first third of the TARA Pacific islands already represents approximately 330 individuals for each genus. The analysis of the samples by 18S barcoding did not provide an assignation beyond the genus level of the colonies collected. Further genetic analyzes are therefore required to determine the genetic structure of the sampled colonies at the species level. Barcoding approaches in corals are notoriously difficult (Shearer and Coffroth 2008), even if mitochondrial diversity may sometimes be used for species delineation ((Flot and Tillier 2007; Paz-García et al. 2016; Johnston et al. 2017). However, in the two genera here studied, previous studies relied on analyses of arrays of dozen microsatellite loci in *Porites* (Baums et al. 2012; Boulay et al. 2014) and *Pocillopora*(Gélin et al. 2017, 2018; Oury 2020; Oury et al. 2022a) or on RAD seq data (Oury et al. 2022b) to reliably identify species.

Metagenomic data were obtained for about one third of the TARA Pacific individuals (Belser et al. 2022) which allowed for a genome wide analysis of these colonies from the first third of the sampled islands (Hume et al. 2022). This study showed that the *Pocillopora* samples in fact belong to five independent genetic lineages or species (SVD1 to SVD5) and that the *Porites* samples corresponded to three lineages (K1-K3). Taking advantage of this genome wide analysis of diversity, we could also identify genomic islands of differentiation among these species (Hume et al. 2022). However, a genome wide SNP analysis is too expensive and resource consuming to be applied even to the full sampling of even the first TARA Pacific 11 islands and could not be used for field conservation purposes.

We therefore developed a single species-diagnostic marker approach to assign the non-sequenced TARA Pacific samples to the species detected in (Hume et al. 2022). This “Divergent Fragment” approach consists in identifying and using a single diagnostic short genomic sequence containing sufficiently divergent linked SNPs to efficiently discriminate the species in large samples.

## Material & Methods

### Sampling

On the 32 islands visited in 2016-2018 by the TARA Pacific expedition, coral colonies were sampled based on morphology, targeting the focal species *Pocillopora meandrina* and *Porites lobata*, as described in Lombard *et al*. (Lombard et al. 2022). On each island, when possible, ten colonies of each species were sampled for each of three different sites. Samples were shipped and stored at Genoscope. Total genomic DNA was extracted from all samples as described in Belser *et al*. (Belser et al. 2022). A third of these samples (3 colonies per site) were sequenced at Genoscope, producing for each colony 1.108 100bp paired end reads of metagenomic data (MetaG dataset) (Belser et al. 2022). The present study focuses on the analysis of the 11 islands from Eastern to Central Pacific of this sampling scheme, which are Las Perlas (I01), Coiba (I02), Malpelo (I03), Easter (I04), Ducie (I05), Gambier (I06), Moorea (I07), Aitutaki (I08), Niue (I09), Samoa (I10), Guam (I15).

### Identification of divergent genomic fragments

The analysis of the MetaG dataset allowed us to genetically identify 5 and 3 species in respectively *Pocillopora* and *Porites* samples (Hume et al. 2022). This result was based on the coalescent analysis of thousands of unlinked genome-wide SNPs and was therefore not easily transposable to the full TARA dataset. The species being identified, the idea was then to track the parts of the genome that diverged the most among them, using this time the full, unfiltered MetaG dataset of linked genome-wide SNPs. The raw MetaG dataset used in Hume *et al*. (2022) was then only filtered on SNP quality, missingness (no missing data) and minimum allele frequency (0,05) using vcftools (Danecek et al. 2011). Genetic differentiation among the species (Weir’s *F*st) was then computed in a 500bp sliding window using vcftools (Danecek et al. 2011). The 1% most divergent 500bp bins were then extracted and sorted to identify the 1 to 2kb long tracts of adjacent bins that contained the highest number of SNPs (this search was limited to the longest genomic contig for *Pocillopora*). The species resolution power of these fragments was tested *in silico* by sNMF clustering (Frichot et al. 2014) of the species assigned, MetaG analyzed individuals using only the SNPs within these genomic fragments. For each coral genus, two to three of these genomic islands of species differentiation were picked as potential targets for amplification.

### PCR set up *in silico*

The first stage for the selection of the divergent fragment was performed *in silico* by extracting from the MetaG dataset the full fasta sequence of the targeted 1 to 2kb fragments. For this, we used the vcfprimers function of vcflib (Garrison et al. 2022), asking to extract the 500bp flanking each SNP included in the genomic island targeted. The full fragment sequence was then reconstructed by assembling these fragments with mafft (Katoh and Standley 2013). All the SNPs contained in the targeted fragment were then placed by hand on this sequence in order to identify conserved tracts that could be used for PCR primers. Potential 25bp primer pairs were then tested using Primer3 (https://primer3.ut.ee/).

### PCR set up *in vitro*

Optimal amplification conditions were tested for each primer pair in a two steps approach. First, a gradient PCR was performed on a single individual to identify the cycling and annealing conditions resulting in the higher possible amplification yield of a single specific fragment. Then, the primer pair conservation was tested by trying to amplify the fragment at these conditions in a sample of one individual per species.

### Nested PCR set up

The amplified divergent fragments were 1,117 and 2,054 bp long for *Pocillopora* and *Porites* respectively. The Illumina sequencing technology we chose allowed for the production of paired ends 250bp reads. We therefore had to set up 500bp long nested PCR along these fragments to prepare the sequencing libraries. 18(+/-2) bp PCR primers were then tested in conserved regions within each of the amplified fragments. 3 and 5 nested PCR products were thus defined for the *Pocillopra* and the *Porites* divergent fragments respectively (Table1 and Figure 1).

**Table 1.**
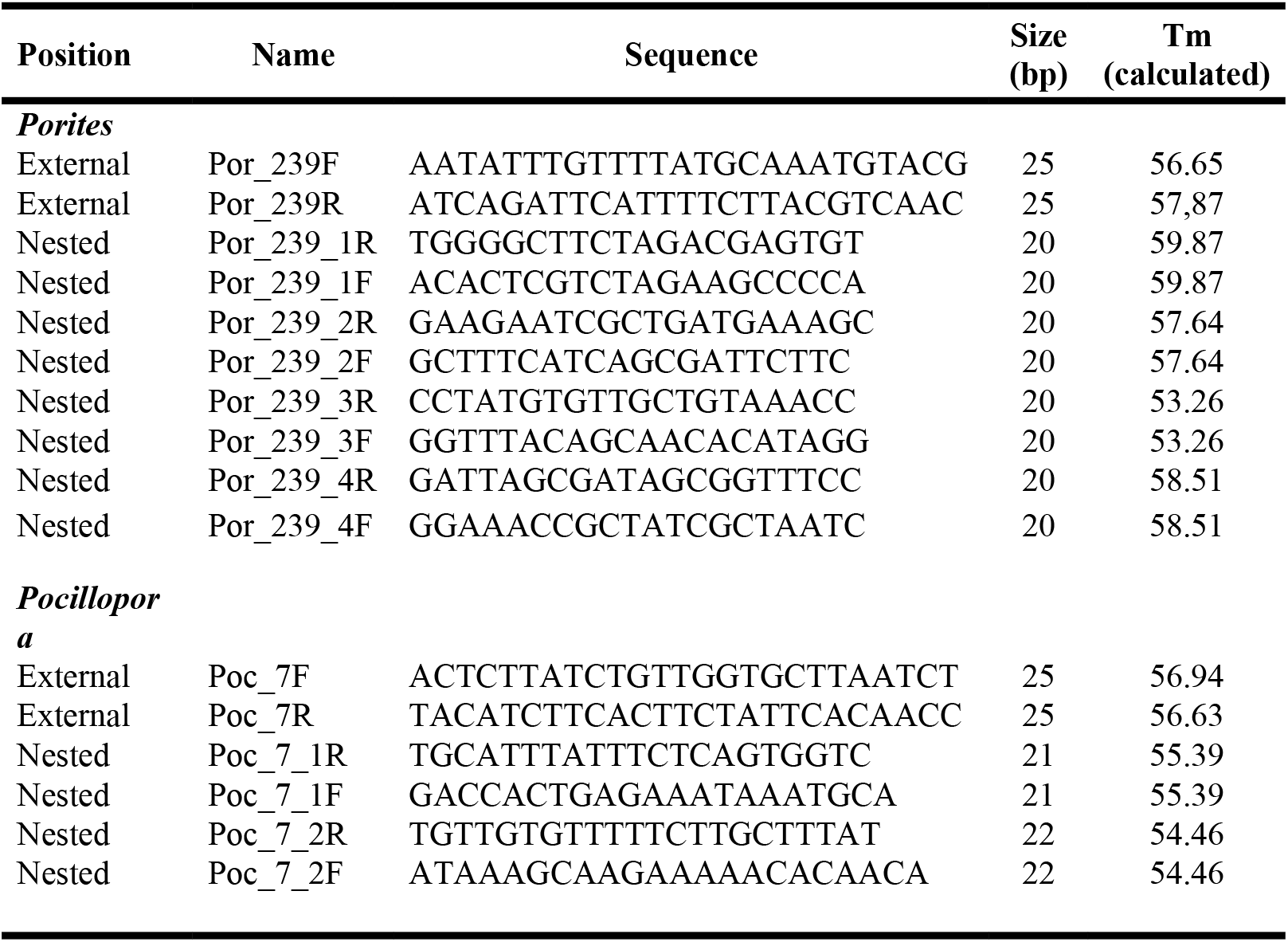
external (PCR) and nested (sequencing) primers.

**Fig. 1.**
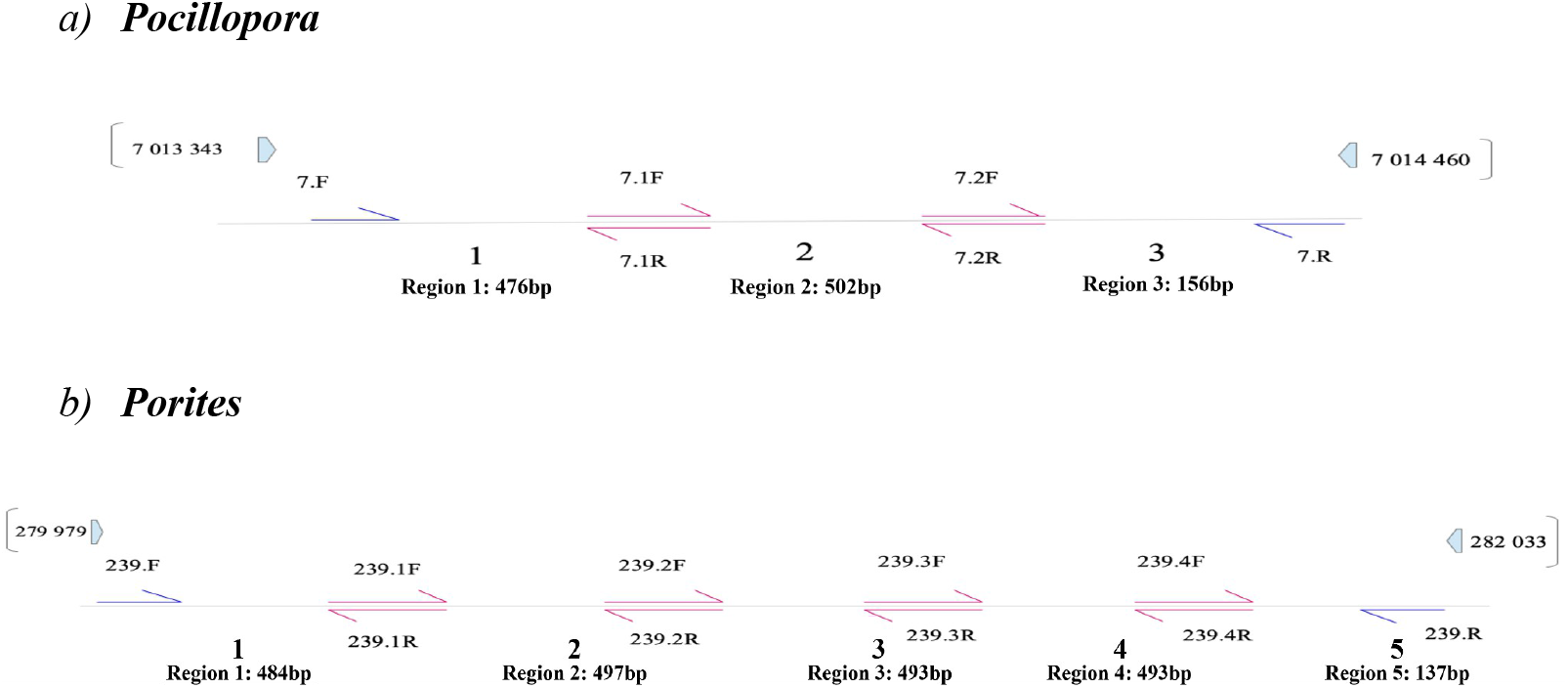
Positions of External and Nested primers and sequences produced for **a)** *Pocillopora* contig 1 from bp 7,013,343 to bp 7,014,460, **b)** *Porites* contig 239 from bp 279,979 to bp 282,033. External PCR primers had to be defined slightly out of the genomic zone tested in silico due to the presence of small indels/SNPs close to the extremities of these zones.

### PCR amplifications

A two step amplification protocol was then performed on each individual total DNA extract. First, the full length divergent fragment was amplified from gDNA samples using the Fidelio® Hot Start PCR kit (OZYA010, Ozyme) with its HF buffer according to the manufacturers’ recommendations and the following PCR programs. 98°C 3 min, 20 cycles of 95°C 30s/58°C 30s/72°C 1 min, and final extension step 72°C 2 min for Pocillopora specific primer pairs. 98°C 3 min, 20 cycles of 95°C 30s/57.5°C 30s/72°C 2 min, and final extension step 72°C 2 min for Porites specific primer pairs. The full length PCR product thus obtained was then purified using paramagnetic beads (AMPure XP – Agencourt A63880), and used as matrix for the nested PCR following the same protocol described above but with a 30s elongation steps at 72°C and 40 cycles (Figure 1).

Nested PCR products were then purified using paramagnetic beads, quantified with a Fluoroskan instrument and validated using a high-throughput microfluidic capillary electrophoresis LabChip GX system (Perkin Elmer, Waltham, MA, USA).

### Library preparation and sequencing

All libraries were prepared using the NEBNext DNA Modules Products and NextFlex DNA barcodes with 100 ng of purified PCR product as input. Purified PCR products were normalized at 2.5 ng / μl. Depending on the size of PCR products, 2 strategies were followed to decrease the number of libraries to prepare. For PCR products of around 500 bp (Poc_7F/Poc_7.1R, Poc_7.1F/Poc_7.2R for *Pocillopora* and Por_239F/Por_239.1R, Por_239.1F/Por_239.2R, Por_239.2F/Por_239.3R, Por_239.3F/Por_239.4R for *Porites*) an equimolar pool of purified PCR products generated from one colony was prepared in order to have a total of 100 ng of amplicons in a total volume of 50 μl. For PCR products around 150 bp (Poc_7.2F/Poc_7R for *Pocillopora* and Por_239.4F/Por_239R for *Porites*) 100 ng of PCR product generated from one colony were used directly as input for the library. 100 ng of pooled PCR products (230 pools for *Pocillopora* and 229 pools for *Porites*) or 100 ng of direct PCR products (229 for *Pocillopora* and 232 for *Porites*) were then end-repaired, A-tailed at the 3’end, and ligated to Illumina compatible adaptors using NEBNext DNA Modules (New England Biolabs, MA, USA) and NextFlex DNA barcodes (BiOO Scientific Corporation, Austin, TX, USA) with Genoscope in-house-developed ‘on beads’ protocol as described in Alberti et al. (Alberti et al. 2017). A liquid handler, the Biomek FX Laboratory Automation Workstation (Beckman Coulter Genomics, Danvers, MA, USA), was used to perform up to 96 reactions in parallel. After two consecutive 1x AMPure XP clean-ups, the ligated products were amplified using Kapa Hifi HotStart NGS library Amplification kit (Kapa Biosystems, Wilmington, MA, USA), followed by 0.6x AMPure XP purification.

All libraries were quantified first by PicoGreen in 96-well plates. Library profiles were assessed using a high throughput microfluidic capillary electrophoresis LabChip GX system (Perkin Elmer, Waltham, MA, USA) and qPCR with the KAPA Library Quantification Kit for Illumina Libraries on an MXPro instrument. Libraries were then sequenced on Illumina sequencer (Illumina, San Diego, CA, USA) in order to obtain an average of 30 to 100 000 of useful paired-end reads.

### Data analysis & species assignation

Sequencing results (fastq files) along with MetaG sequences from individuals of known species assignation (at least 2 per species) were aligned on the reference sequence of the divergent fragment used for PCRsetup using BWA-mem v0.7.15 (Li and Durbin 2009). The obtained sam files were then translated into bam format by samtools v1.10.2 (Li et al. 2009), and the bam files were subsequently sorted and duplicate reads were marked by sambamba (Tarasov et al. 2015). Variable positions were then called by samtools mpileup (Danecek et al. 2021) from all the obtained bam files for each genus. The obtained vcf files were then further filtered for SNP only (no indels), missingness (no missing data) and minimum allele frequency (0,05) with vcftools (Danecek et al. 2011). This file was then converted to plink format with vcftools, and a sNMF hierarchical clustering analysis of this dataset was then performed using the LEA R package (Frichot et al. 2014).

## Results/Discussion

### *In silico* species diagnostic validation

To test which divergent genomic fragment could be species diagnostic, the SNPs contained within each putative genomic fragment were extracted from the whole genomic datasets on which the species delineation was performed (Hume et al. 2022), and a species assignation was performed on these SNPs only. The results of these assignations for two candidate fragments (one from *Pocillopora* contig 1, the other from *Porites* contig 239), are compared to the genome wide SNP based assignations from the same individuals in Tables 2a and 2b for *Pocillopora* and *Porites* respectively. As can be seen, in both genera, for these two fragments, the two assignations were concordant for more than 97% of the samples (3 colonies over 105 tested were mistyped in *Pocillopora* and 2 in 109 for *Porites*). Considering that in each genus, one of the very few mistyped colonies was detected as an hybrid in the genome wide SNP analysis, this result clearly validated these two genomic fragments as potential species diagnostic divergent fragments.

**Table 2.** *in slico* Divergent fragment validation test. These tables give the species assignation results based on either the full set of genowide wide SNPs (from Hume at al 2022) in columns or from the putative divergent fragment SNPs in a) *Pocillopora* (SNPs from contig 1, from bp 7013501 to bp 7014500), b) *Porites* (SNPs from contig 239, from bp 280001 to bp 282000)

|  |  | Genome wide SNPs |  |  |  |  |  | Total |
| --- | --- | --- | --- | --- | --- | --- | --- | --- |
|  |  | SVD1 | SVD2 | SVD2/SVD3 | SVD3 | SVD4 | SVD5 |  |
| Fragment | SVD1 | 12 | - | - | - | - | - | 12 |
|  | SVD2 | - | 18 | 1 | - | 2 | - | 21 |
|  | SVD3 | - | - | - | 18 | - | - | 18 |
|  | SVD4 | - | - | - | - | 34 | - | 34 |
|  | SVD5 | - | - | - | - | - | 20 | 20 |
| Total |  | 12 | 18 | 1 | 18 | 36 | 20 | 105 |

b) *Porites*
|  |  | Genome wide SNPs |  |  |  | Total |
| --- | --- | --- | --- | --- | --- | --- |
|  |  | K1 | K2 | K2/K3 | K3 |  |
| Fragment | K1 | 23 | - | 1 | - | 24 |
|  | K2 | - | 35 | - | - | 35 |
|  | K3 | - | 1 | - | 49 | 50 |
| Total |  | 23 | 36 | 1 | 49 | 109 |

### Sequencing results

The careful *in silico* selection of PCR primers allowed for successful amplification of the first divergent fragments tested. Sequences were obtained for 225 and 241 individual colonies respectively. After SNP calling and quality filters, the final vcf file contained 26 SNPs in the 1117kb long divergent fragment in *Pocillopora* and 59 SNPs in the 2054kb long *Porites* divergent genomic fragment. This resulted in 25 unique genotypes among 235 *Pocillopora* individuals and 28 unique genotypes among 241 *Porites* individuals.

### Species assignation

As shown in Figure 2a, sNMF analyses stated that the optimal number of ancestral populations (*i*.*e*. genetic clusters) to assign the individuals to was K=5 in *Pocillopora*. Each pair of reference individuals was unambiguously assigned to a different cluster, which allowed to assign to each of these clusters one and only one species identified by genome wide analyses. As shown in Table 3, among the 235 sequenced individuals, 219 were unambiguously assigned to only one of the five clusters/species (Assignation Q score > 0.80 in one cluster). 16 individuals were detected as hybrids by sNMF (Assignation Q score < 0.80 in all clusters, but > 0.25 in some), and this hybridization was confirmed by the polymorphic positions in their genotypes (Table 4). In *Porites*, the sNMF analysis of the divergent fragment genotypes pointed to an optimal number of ancestral populations of K=4 (Figure 2b), one more cluster than the three species detected by genome wide analysis (Hume et al. 2022). However, the reference individuals were all unambiguously assigned to only three clusters (data not shown). We therefore forced a K=3 assignation in sNMF, and all the individuals of the previous fourth cluster were then assigned to the same cluster as the K3 reference individuals. The K=3 assignation results are shown in Table 5. As in *Pocillopora*, 13 hybrid individuals could be detected by both sNMF and genotype compositions (Table 6).

**Table 3.** Species assignations form the divergent fragment and genome wide SNPs in *Pocillopora*. For each island, the number of individual colonies assigned to a given species is given in italic for the genome wide SNP analysis (from Hume *et al*. 2022), and in bold for the divergent fragment analysis

| Species assignment | Island |  |  |  |  |  |  |  |  |  |  | Total |
| --- | --- | --- | --- | --- | --- | --- | --- | --- | --- | --- | --- | --- |
|  | I01 | I02 | I03 | I04 | I05 | I06 | I07 | I08 | I09 | I10 | I15 |  |
| **SVD1** | - | - | - | - | *4 / 8* | *7 / 11* | *1 / 1* | - | - | - | - | *12 / 20* |
| **SVD2** |  |  |  |  | *- / 1* | *- / 1* | *1 / 5* | *4 / 14* | *3 / 8* | *7 / 16* | *3 / 8* | *18 / 53* |
| **SVD2/SVD3** | - | - | - | - | - | - | - | - | *1 / -* | - | - | *1 / -* |
| **SVD2/SVD4** | - | *- / 2* | - | - | - | *- / 1* | - | - | - | - | - | *- / 3* |
| **SVD2/SVD5** | - | - | - | - | *- / 1* | *- / 1* | - | *- / 3* | - | *- / 2* | - | *- / 7* |
| **SVD3** | - | - | - | - | - | - | - | *5 / 1* | *5 / 14* | *2 / 5* | *6 / 14* | *18 / 34* |
| **SVD3/SVD5** | - | - | - | - | - | - | *- / 1* | - | - | - | - | *- / 1* |
| **SVD4** | *11/24* | *10/25* | *10/ -* | - | *3 / 5* | *2 / 2* | - | - | - | - | - | *36/56* |
| **SVD4/SVD2/SVD5** | *- / 1* | - | - | - | *- / 1* | - | - | - | - | - | - | */ 2* |
| **SVD5** | - | - | - | *12/23* | *2 / 7* | *- / 5* | *7 / 17* | */ 4* | - | - | - | *21/56* |
| **SVD5/SVD2/SVD4** | - | - | - | *- / 1* | - | - | - | - | - | - | - | *- / 1* |
| **SVD5/SVD4** | - | - | - | *- / 2* | - | - | - | - | - | - | - | *- / 2* |
| **Total** | *11/25* | *10/27* | *10/ -* | *12/26* | *9/23* | *9/21* | *9/24* | *9/22* | *9/22* | *9/23* | *9/22* | *106/235* |

**Table 4.**
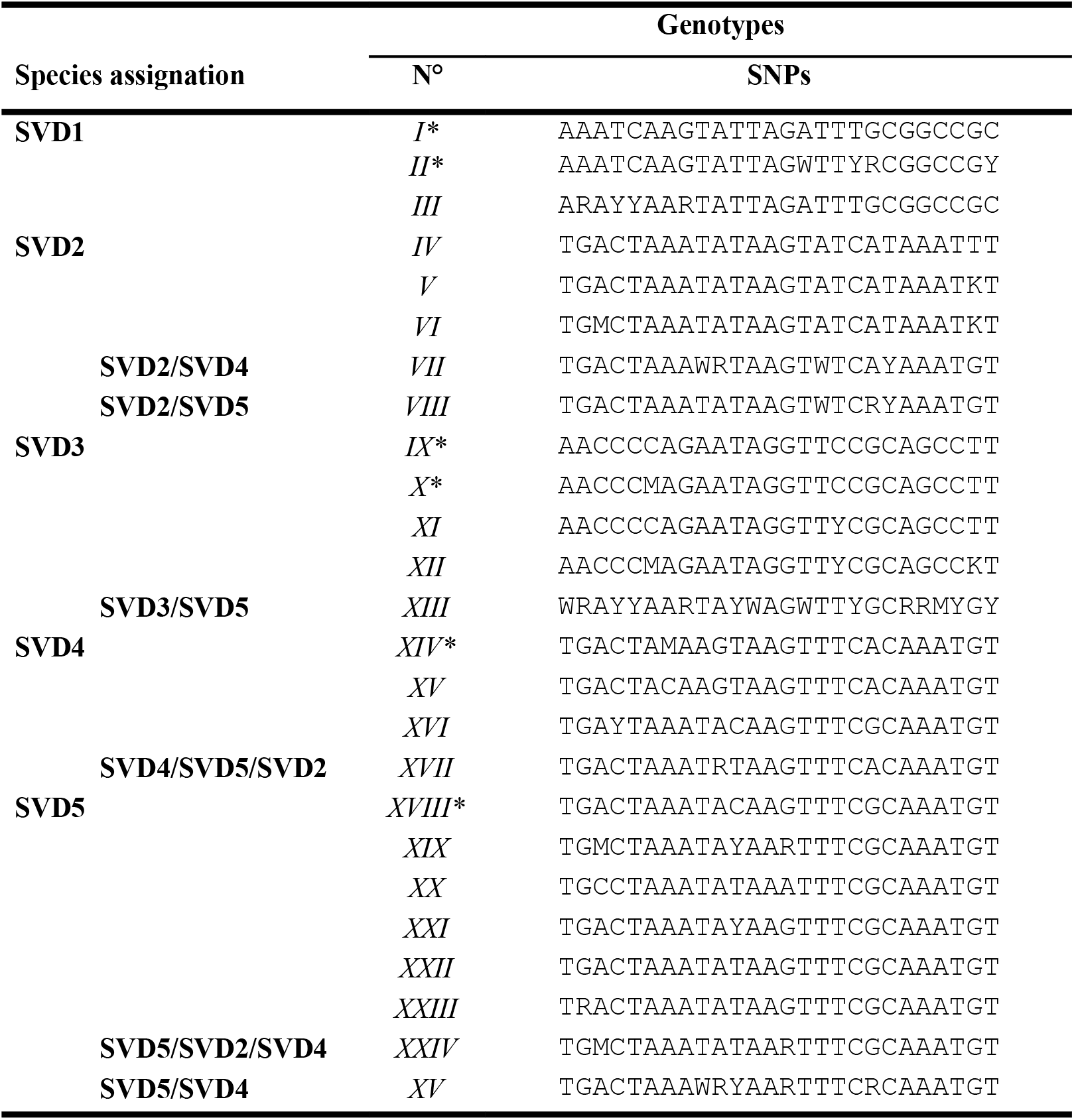
Genotypes obtained in *Pocillopora* and their species assignation. Each genotype is designated by a number, an * indicates this genotype was found in at least one reference individual.

**Table 5.** Species assignations form the divergent fragment and genome wide SNPs in *Porites*. For each island, the number of individual colonies assigned to a given species is given in italic for the genome wide SNP analysis (from Hume et al. 2022), and in bold for the divergent fragment analysis

| Species assignment | Island |  |  |  |  |  |  |  |  |  |  | Total |
| --- | --- | --- | --- | --- | --- | --- | --- | --- | --- | --- | --- | --- |
|  | I01 | I02 | I03 | I04 | I05 | I06 | I07 | I08 | I09 | I10 | I15 |  |
| **K1** | *9 / 22* | *2 / 11* | *12 / 9* | - | - | - / 1 | - | - | - | - | - | *23 / 43* |
| **K2** | - | **7 / 8** | - / 3 | - | *2 / 1* | *2 / 4* | *7 / 13* | *7 / 3* | *2 / -* | *9 / 23* | *4 / 11* | *36 / 66* |
| **K2/K3** | - | - / 4 | - / 3 | - | - | - / 2 | - / 2 | - | - | - | *1 / -* | *1 / 11* |
| **K3** | - | - | - / 1 | *12 / 29* | *7 / 22* | *7 / 14* | *6 / 2* | *6 / 19* | *7 / 22* | - / 1 | *5 / 9* | *49 / 119* |
| **K4 ? (K1/K3/K3)** | - | - | - | - | - | - | - | - | - | - | - / 2 | - / 2 |
| **Total** | *9 / 22* | *9 / 23* | *12 / 16* | *12 / 29* | *9 / 23* | *9 / 21* | *13 / 17* | *9 / 22* | *9 / 22* | *9 / 24* | *9 / 22* | *109 / 241* |

**Table 6.**
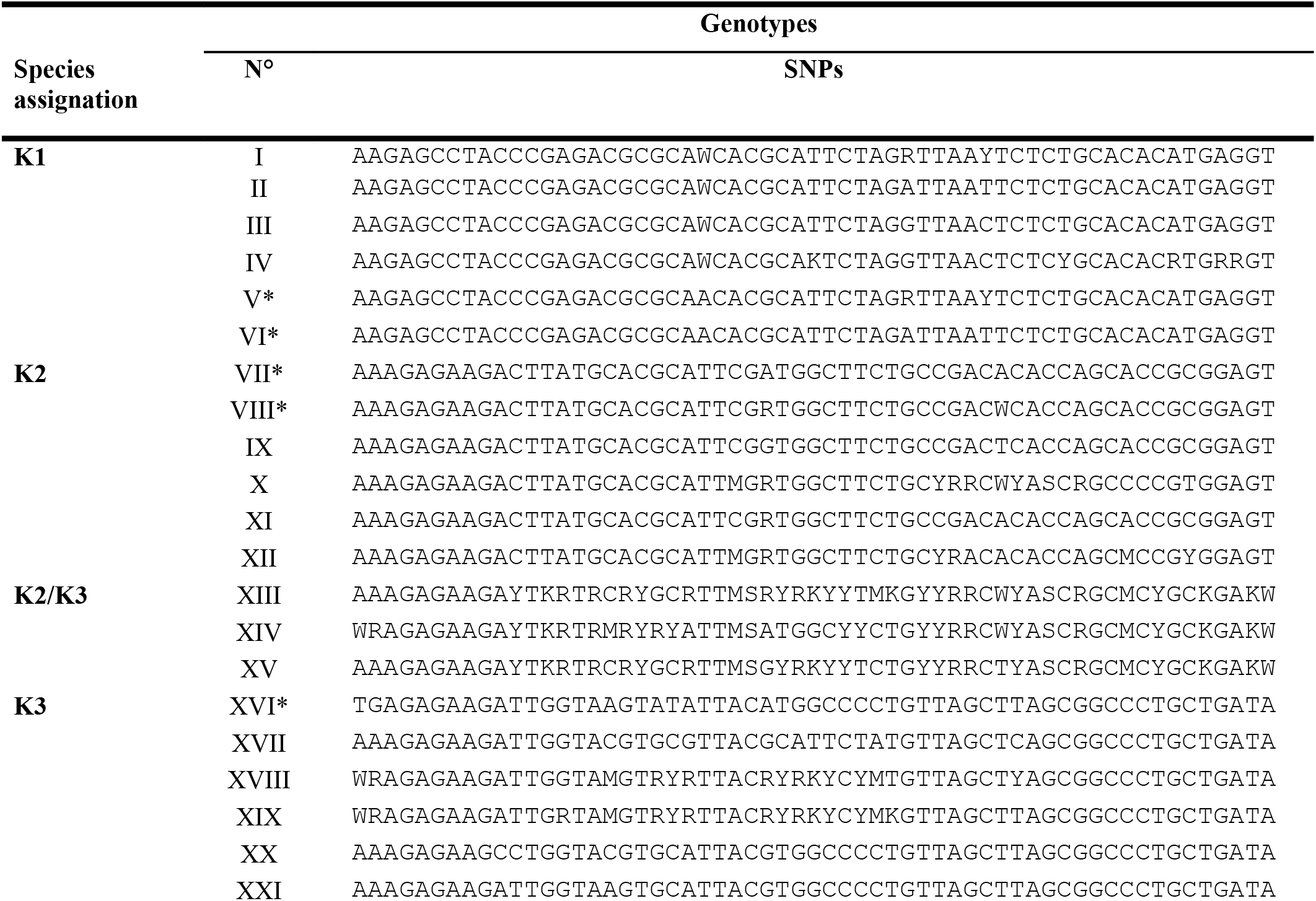
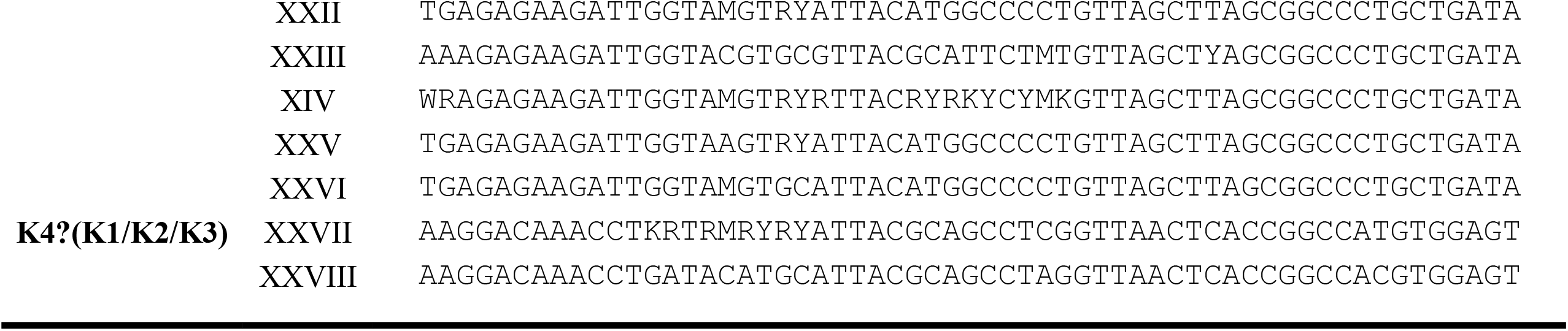
Genotypes obtained in *Porites* and their species assignation. Each genotype is designated by a number, an * indicates this genotype was found in at least one reference individual.

**Fig. 2.**
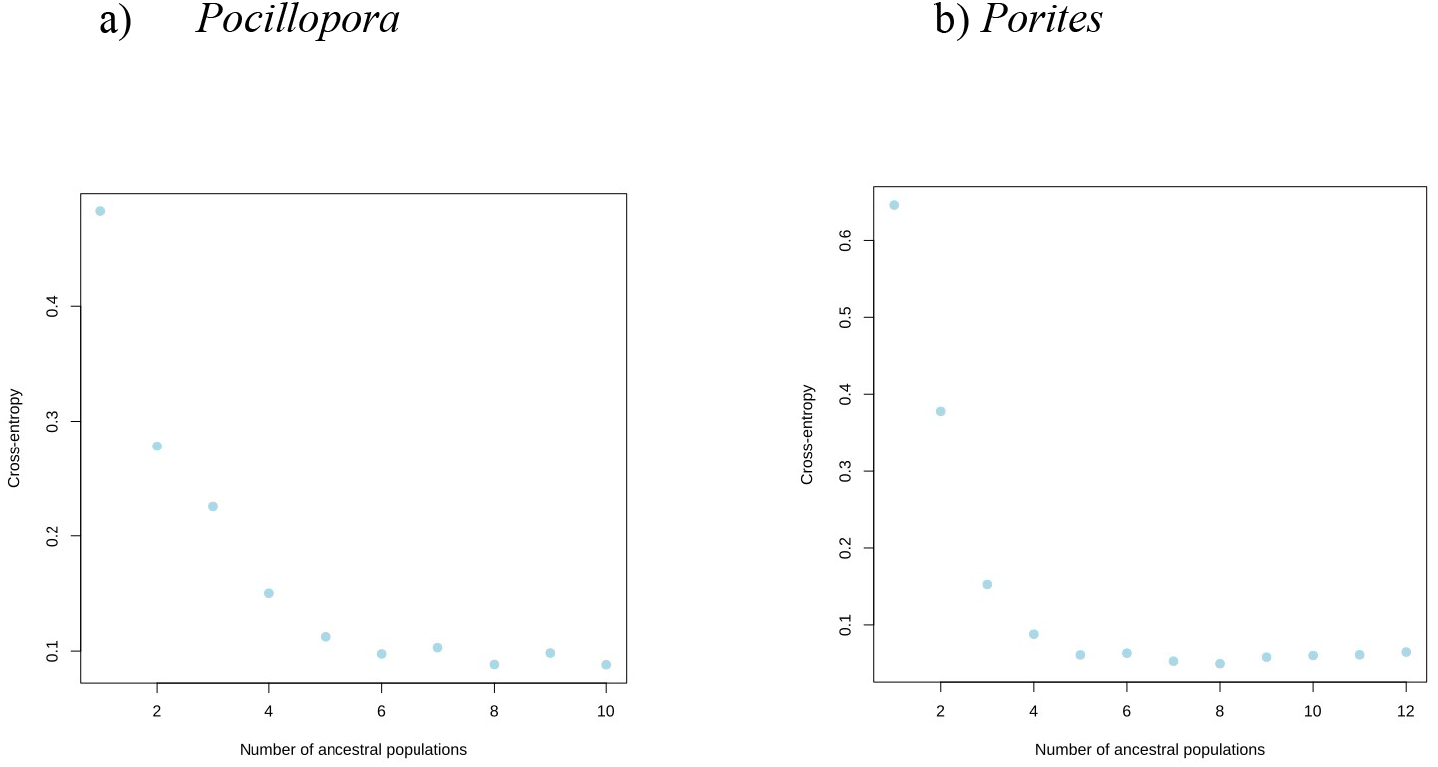
sNMF Entropy graphs. These graphs identify the optimal number of ancestral populations (*i*.*e*. clusters) to which the individual sequences are assigned. a) *Pocillopora*. The optimal number of ancestral populations was K=5, which corresponded to the number of species identified by genome wide analysis; b) *Porites*. Here the optimal number of ancestral populations was K=4, when three species were detected by genome wide analysis. However, the third and fourth clusters were totally merged when forcing a K=3 clustering.

The Tables 3 and 5 also show the assignation results previously obtained by genome wide analysis (Hume et al. 2022). Though using a different and more numerous set of individuals, the biogeographic distribution of the species obtained with the divergent fragment is highly comparable with these previous results. The accrued sample size in the divergent fragment analysis also probably allowed for the detection of a fourth species in *Porites*, that could correspond to the two individuals from I15 that could not be clearly assigned to any of the three clusters. The genome wide analysis of the full 32 islands samples should confirm the existence of this fourth species, if it becomes more frequent in Western Pacific.

This work made it possible to assign individual coral colonies to species in two genera of corals (*Pocillopora* and *Porites*) collected from 11 islands from eastern to central Pacific during the Tara Pacific expedition, by only amplifying and sequencing a single short genomic sequence of less than 2kb. This simple species assignation tool has thus the same resolution as the genome wide analyses (Hume et al. 2022, Oury et al. 2022b) or as the microsatellite arrays (Boulay et al. 2014; Gélin et al. 2017) which were used so far in these genera. It should however be noted that the identification of these diagnostic divergent fragments stemmed from a previous full-fledged genome wide analysis. The diagnostic value of these fragments outside of the geographic zone here sampled still also needs to be tested. The upcoming analysis of the full set of TARA Pacific 32 islands will show if these divergent fragments are still diagnostic at the full Pacific basin scale.

The taxonomic information provided here has already proved an essential asset for crossover analyzes of phenotypic in this geographical area, allowing for a better understanding of the impact of environmental variation on coral reefs.

## Acknowledgements

We are keen to thank the commitment of the people and the following institutions for their financial and scientific support that made Tara Pacific expedition possible: CNRS, PSL, CSM, EPHE, Genoscope/CEA, Inserm, Université Côte d’Azur, ANR, Agnès b., UNESCO-IOC, the Veolia Environment Foundation, Région Bretagne, Billerudkorsnas, Amerisource Bergen Company, Lorient Agglomeration, Smilewave, Oceans by Disney, the Prince Albert II de Monaco Foundation, L’Oréal, Biotherm, France Collectivités, Fonds Français pour l’Environnement Mondial (FFEM), the Ministère des Affaires Européennes et Etrangères, the Museum National d’Histoire Naturelle, Etienne BOURGOIS, the Tara Ocean Foundation’s teams and crew. Tara Pacific would not exist without the continuous support of the participating institutes. The authors also particularly thank Serge Planes, Denis Allemand and the Tara Pacific consortium.

The acknowledgements to local people and authorities who made this study possible can be found in the annex.

DF work was supported by the French Government (National Research Agency, ANR) through the grant “Coralgene” ANR-17-CE02-0020

The Tara Pacific expedition would not have been possible without the participation and commitment of over 200 scientists, sailors, artists and citizens (see https://zenodo.org/record/3777760#.YfEEsfXMLjB).

## Annex

## Acknowledgements

**Colombia :**

We are keen to thank the commitment of the people of Colombia and the following local institutions for their administrative and scientific support that made this singular expedition possible : the Prime Minister Office and their teams; the Ministry of Environmen and National Natural Parks of Colombia with Guillermo Alberto Santos Ceballos, Edna Carolina Jarro Fajardo, and their teams; la Fundacion Malpelo y Otros Ecosistemas Marinos with their Executive Director and Founder Sandra Bessudo, with Felipe Orlando Ladino and their teams; el Santuario de Fauna y Flora Malpelo and their teams; la Direccion General Maritima with Ivan Fernando Castro Mercado and their teams; The Pacific Territorial Directorate and their teams; the Colombian biodiversity information system and their teams; and all other locally supports.

**Panama :**

We are keen to thank the commitment of the people of Panama and the following local institutions for their administrative and scientific support that made this singular expedition possible : The Ministry of Environment of Panama with Samuel Valdès Diaz, Lissette Trejos, Patricia Hernandez and their teams; the Smithsonian Tropical Research Institute with Rachel Page, Zurenayka Alain, Juan Mate and their teams; the Head of MPA at Coiba Park Didiel Nunez and their teams; and all other locally supports.

**Pitcairn Islands :**

We are keen to thank the commitment of the people of Pitcairn Islands and the following local institutions for their administrative and scientific support that made this singular expedition possible : the Government of Pitcairn Islands with Melva Evans and their teams; the Office of the Mayor and Council for Pitcairn, Henderson, Ducie and Oeno Islands with Shawn Christian and their teams; the Environmental, Conservation & Natural Resources Division with her Manager Michele Christian, and their teams; the Police & Immigration Officer with Christian Brenda, and their teams; and all other locally supports.

**French Polynesia :**

We are keen to thank the commitment of the people of French Polynesia and the following local institutions for their administrative and scientific support that made this singular expedition possible : Le Haut-commissaire de la République en Polynésie française and their teams; The City of Papeete with its Mayor, its Port Commander, and their teams; le Centre de Recherches Insulaires et Observatoire de l’Environnement (CRIOBE) and their teams; and all other locally supports.

**Cook Islands :**

We are keen to thank the commitment of the people of the Cook Islands and the following local institutions for their administrative and scientific support that made this singular expedition possible : the Government of the Cook Islands and their teams; the Office of the Prime Minister with Tina Samson, Elizabeth Wright-Koteka from the

Cook Islands Research Committee, and their teams; the Ministry of Marine Resources with its Director Dorothy Solomona, and their teams; the National Environment Service with its Director Joseph Brider, Elizabeth Munro, Bobby Bishop, and their teams; the City of Aitutaki, its mayor, and their teams; the Aitutaki Marine Research Station with its station manager Richard Story, and their teams; the Cawthron Institute with Lesley Rhodes, Kirsty Smith, and their teams; and all other locally supports.

**Niue :**

We are keen to thank the commitment of the people of Niue and the following local institutions for their administrative and scientific support that made this singular expedition possible : the Government of Niue with Richard Hipa, James Tatafu, Aldric Hipa, and their teams; the Office of External Affairs of the Government of Niue with its Director Emi Hipa, and their teams; the Cabinet and Parliamentary Services with its Director Christine External; the Ministry of Infrastructure and its Department of Transport, with Lynsey Talagi, and their teams; the Ministry of Natural Resources and its Director-General Josie Tamate, and their teams; the Department of Agriculture, Forestry and Fisheries at the Ministry of Natural Resources with its Director Brendon Pasisi, and their teams; the Department of Environment of the Ministry of Natural Resources and its Director Sauni Tongatule, and their teams; the Pacific Community with its Deputy Director-General Cameroun Diver, Coral Pasisi, and their teams; and all other locally supports.

**Samoa Islands :**

We are keen to thank the commitment of the people of Samoa Islands and the following local institutions for their administrative and scientific support that made this singular expedition possible : the Government of the Independent State of Samoa, and their teams; the Ministry of Foreign Affairs and Trade with its CEO Peseta Noumea Simi, its Acting CEO Leroy E. Hunkin-Mamae, Sapeti Titiii, and their teams; the Ministry of Natural Resources and Environment with its Principal Marine Conservation Officer Maria R. Satoa Peni, and their teams; the Ministry of Agriculture and Fisheries with its Assistant CEO Magele Etuati Ropeti, and their teams; and all other locally supports.

**Guam (US) :**

We are keen to thank the commitment of the people of Guam and the following local institutions for their administrative and scientific support that made this singular expedition possible : the Department of Agriculture Division of Aquatic and Wildlife Resources (DAWR), its Director Matthew L.G. Sablan, its Assistant Chief Jay Gutierrez, and their teams; the Division of Management Authority Branch of Permits, the Senior Biologist Anna Barry, and their teams; the U.S. Fish and Wildlife Service – Office of Law Enforcement, its Wildlife Inspector Arthur T. Taimanglo, and their teams; and all other locally supports.

## References

Alberti A, Poulain J, Engelen S, et al (2017) Viral to metazoan marine plankton nucleotide sequences from the Tara Oceans expedition. Sci Data 4:170093. https://doi.org/10.1038/sdata.2017.93

Baums IB, Boulay JN, Polato NR, Hellberg ME (2012) No gene flow across the Eastern Pacific Barrier in the reef-building coral Porites lobata. Molecular Ecology n/a-n/a. https://doi.org/10.1111/j.1365-294X.2012.05733.x

Belser C, Poulain J, Labadie K, et al (2022) Integrative omics framework for characterization of coral reef ecosystems from the Tara Pacific expedition. DOI:https://doi.org/10.48550/arXiv.2207.02475

Boulay JN, Hellberg ME, Cortés J, Baums IB (2014) Unrecognized coral species diversity masks differences in functional ecology. Proc R Soc B 281:20131580. https://doi.org/10.1098/rspb.2013.1580

Chen P-Y, Chen C-C, Chu L, McCarl B (2015) Evaluating the economic damage of climate change on global coral reefs. Global Environmental Change 30:12–20. https://doi.org/10.1016/j.gloenvcha.2014.10.011

Danecek P, Auton A, Abecasis G, et al (2011) The variant call format and VCFtools. Bioinformatics 27:2156–2158

Danecek P, Bonfield JK, Liddle J, et al (2021) Twelve years of SAMtools and BCFtools. GigaScience 10:giab008. https://doi.org/10.1093/gigascience/giab008

Flot JF, Tillier S (2007) The mitochondrial genome of Pocillopora (Cnidaria: Scleractinia) contains two variable regions: the putative D-loop and a novel ORF of unknown function. Gene 401:80–87

Frichot E, Mathieu F, Trouillon T, et al (2014) Fast and Efficient Estimation of Individual Ancestry Coefficients. Genetics 196:973–983. https://doi.org/10.1534/genetics.113.160572

Garrison E, Kronenberg ZN, Dawson ET, et al (2022) A spectrum of free software tools for processing the VCF variant call format: vcflib, bio-vcf, cyvcf2, hts-nim and slivar. PLOS Computational Biology 18:e1009123. https://doi.org/10.1371/journal.pcbi.1009123

Gélin P, Fauvelot C, Bigot L, et al (2018) From population connectivity to the art of striping Russian dolls: the lessons from Pocillopora corals. Ecology and Evolution 8:1411–1426. https://doi.org/10.1002/ece3.3747

Gélin P, Postaire B, Fauvelot C, Magalon H (2017) Reevaluating species number, distribution and endemism of the coral genus Pocillopora Lamarck, 1816 using species delimitation methods and microsatellites. Molecular Phylogenetics and Evolution 109:430–446. https://doi.org/10.1016/j.ympev.2017.01.018

Hume BC, Voolstra CR, Armstrong E, et al (2022) Disparate patterns of genetic divergence in three widespread corals across a pan-Pacific environmental gradient highlights species-specific adaptation trajectories. bioRxiv 2022.10.13.512013

Johnston EC, Forsman ZH, Flot J-F, et al (2017) A genomic glance through the fog of plasticity and diversification in Pocillopora. Scientific Reports 7:. https://doi.org/10.1038/s41598-017-06085-3

Katoh K, Standley DM (2013) MAFFT Multiple Sequence Alignment Software Version 7: Improvements in Performance and Usability. Molecular Biology and Evolution 30:772–780. https://doi.org/10.1093/molbev/mst010

Li H, Durbin R (2009) Fast and accurate short read alignment with Burrows–Wheeler transform. bioinformatics 25:1754–1760

Li H, Handsaker B, Wysoker A, et al (2009) The Sequence Alignment/Map format and SAMtools. Bioinformatics 25:2078–2079. https://doi.org/10.1093/bioinformatics/btp352

Lombard F, Bourdin G, Pesant S, et al (2022) Open science resources from the Tara Pacific expedition across coral reef and surface ocean ecosystems. bioRxiv 2022.05.25.493210. https://doi.org/10.1101/2022.05.25.493210

Masson-Delmotte V, Intergovernmental Panel on Climate Change, WMO, United Nations Environment Programme (2018) Global warming of 1.5 oC: an IPCC special report on the impacts of global warming of 1.5 °C above pre-industrial levels and related global greenhouse gas emission pathways, in the context of strengthening the global response to the threat of climate change, sustainable development, and efforts to eradicate poverty: summary for policy-makers. Intergovernmental Panel on Climate Change, Geneva

Oury N (2020) Cryptic species and genetic connectivity among populations of the coral Pocillopora damicornis (Scleractinia) in the tropical southwestern Pacific. Marine Biology 16

Oury N, Gélin P, Rajaonarivelo M, Magalon H (2022a) Exploring the Pocillopora cryptic diversity: a new genetic lineage in the Western Indian Ocean or remnants from an ancient one? Mar Biodivers 52:5. https://doi.org/10.1007/s12526-021-01246-0

Oury N, Noël C, Mona S, et al (2022b) From Genomics to Integrative Taxonomy? The Case Study of Pocillopora Corals. 2022.10.04.510617

Paz-García D, Galván-Tirado C, Alvarado J, et al (2016) Variation in the whole mitogenome of reef-building Porites corals. Conservation Genetics Resources 8:123–127. https://doi.org/10.1007/s12686-016-0527-x

Planes S, Allemand D, Agostini S, et al (2019) The Tara Pacific expedition—A pan-ecosystemic approach of the “-omics” complexity of coral reef holobionts across the Pacific Ocean. PLoS Biol 17:e3000483. https://doi.org/10.1371/journal.pbio.3000483

Shearer TL, Coffroth MA (2008) Barcoding corals: limited by interspecific divergence, not intraspecific variation. Molecular Ecology Resources 8:247–255

Tarasov A, Vilella AJ, Cuppen E, et al (2015) Sambamba: fast processing of NGS alignment formats. Bioinformatics 31:2032–2034. https://doi.org/10.1093/bioinformatics/btv098

